# Combinatorial T cell engineering eliminates on-target off-tumor toxicity of CD229 CAR T cells while maintaining anti-tumor activity

**DOI:** 10.1101/2021.12.06.471279

**Authors:** Erica R. Vander Mause, Jillian M. Baker, Kenneth A. Dietze, Sabarinath V. Radhakrishnan, Thierry Iraguha, Patricia Davis, Jens Panse, James E. Marvin, Michael L. Olson, Mary Steinbach, David P. Ng, Carol S. Lim, Djordje Atanackovic, Tim Luetkens

**Affiliations:** Department of Microbiology and Immunology, University of Maryland School of Medicine, Baltimore, MD, United States; Hematology and Hematologic Malignancies, Huntsman Cancer Institute, University of Utah, Salt Lake City, UT, United States; Department of Pharmaceutics & Pharmaceutical Chemistry, University of Utah, Salt Lake City, UT, United States; Division of Hematology and Oncology, Medical College of Wisconsin, Milwaukee, USA; Division of Experimental and Clinical Pathology, ARUP Laboratories, Salt Lake City, USA; Department of Pathology, University of Utah, Salt Lake City, UT, United States; Department of Oncology, Hematology, Hemostaseology, and Stem Cell Transplantation, University Hospital RWTH Aachen, Aachen, Germany; Huntsman Cancer Institute, University of Utah, Salt Lake City, UT, United States; Department of Medicine and Transplant/Cell Therapy Program, University of Maryland School of Medicine and Marlene and Stewart Greenebaum Comprehensive Cancer Center, Baltimore, MD, United States

## Abstract

T cells expressing chimeric antigen receptors have shown remarkable therapeutic activity against different types of cancer. However, their wider use has been hampered by the potential for life-threatening toxicities due to the unintended targeting of healthy cells expressing low levels of the targeted antigen. We have now developed an affinity-tuning approach for the generation of minimally modified, low-affinity antibody variants derived from existing high-affinity antibodies. Using this approach, we engineered low affinity variants of the fully human CD229-specific antibody 2D3. Parental 2D3 originally efficiently targeted multiple myeloma cells but also healthy T cells expressing low levels of CD229. We demonstrate that CAR T cells based on a low affinity variant of 2D3, engineered to also express CJUN to increase CAR T cell expansion, maintain the parental antibody’s anti-tumor activity but lack its targeting of healthy T cells *in vitro* and *in vivo*. In addition, we found that low affinity CD229 CAR T cells show reduced trogocytosis potentially augmenting CAR T cell persistence. The fast off-rate CAR produced using our affinity tuning approach eliminates a key liability of CD229 CAR T cells and paves the way for the effective and safe treatment of patients with multiple myeloma and other lymphoid malignancies.

**One sentence summary:** Rational T cell engineering yields low affinity CD229 CAR T cells overexpressing CJUN, which maintain the parental cells’ anti-tumor activity but eliminate killing of healthy T cells, increasing CAR T cell expansion, and decreasing trogocytosis.

## MAIN TEXT

T cells expressing chimeric antigen receptors using antibodies to target cancer-associated surface antigens have been shown to be highly effective against several hematologic malignancies, including B cell lymphoma^1^ and multiple myeloma^2, 3^. However, their extraordinary cytotoxic activity poses new challenges, such as the unintended killing of healthy tissues expressing the targeted antigen, despite often at substantially lower levels^4^. In the case of the widely used CD19 CAR T cells, this on-target off-tumor toxicity results in the elimination of healthy B cells^5, 6^ and various other CAR T cell approaches have resulted in life-threatening toxicities and even patient deaths due to the targeting of healthy tissues^7–9^. It has been shown previously that CAR T cells exert potent anti-tumor activity across a wide range of affinities^10–12^ and many CAR T cell strategies currently in clinical use likely exceed the required affinity threshold. As a consequence, several low affinity antibodies have been developed for a number of cancer targets to increase cancer selectivity as well as CAR T cell persistence and function^13–16^. However, none of these binding domains were derived from existing and extensively tested high affinity antibodies already in clinical use and had to again undergo rigorous preclinical evaluation with the risk for substantial liabilities, such as off-target reactivity and unstable epitopes.

We have now developed an affinity-tuning platform for the generation of low-affinity antibody variants derived from existing high-affinity antibodies. We demonstrate that CAR T cells based on antibody variants developed using this approach show increased selectivity for tumor cells, increased expansion, maintained anti-tumor activity *in vitro* and *in vivo*, and reduced trogocytosis, the stripping of target antigen from tumor cells by CAR T cells, potentially augmenting their persistence *in vivo*. We hypothesize that approaches enabling the systematic reduction in antibody affinity of existing CAR binding domains will become an important tool in the development of more effective CAR T cell approaches.

## RESULTS

### Generation of CD229 antibody variants to increase CAR T cell selectivity

CAR T cells targeting BCMA, an antigen otherwise exclusively expressed on plasma cells, have been approved for the treatment of multiple myeloma (MM)^17, 18^. However, most patients relapse within the first year ^2^ potentially due to incomplete targeting of MM-propagating cells in the memory B cell pool^19, 20^. Recently, an alternative CAR T cell approach based on the phage-display derived, fully human anti-CD229 antibody 2D3 was developed showing targeting not only of terminally differentiated MM plasma cells but also of MM-propagating cells ^21^. While CD229 CAR T cells indeed show efficient targeting of MM cells (Fig. 1A, Suppl. Fig. 1), they also target healthy T cells (Fig. 1B), indicating the potential for substantial toxicities. Analyzing expression of CD229 on MM cells and normal T cells from patients with relapsed-refractory MM using flow cytometry, we found that T cells express significantly lower levels of CD229 than MM cells (Fig. 1C). Affinity-tuning of CAR binding domains has previously been shown to reduce targeting of cells expressing lower levels of the targeted antigen (Fig. 1D). Compared to other commonly used CAR binding domains, such as the CD19-specific antibody FMC63^22^, the affinity of wildtype 2D3 is already relatively low at 476nM (Fig. 1E). Considering the high specificity and extensive preclinical characterization of 2D3, as well as the established anti-tumor activity and functionality of 2D3-based CAR T cells^21^, we developed an affinity tuning platform to generate low affinity variants of the 2D3 binding domain by comprehensively mutating heavy and light CDR3 regions (Fig. 1F) in combination with high-throughput screening and antibody characterization assays. We generated 305 single amino acid substitution variants of 2D3 (Fig. 1G), with the goal to substantially reduce 2D3 affinity, while maintaining the recognized CD229 epitope, 2D3 specificity, expansion, and anti-tumor activity.

**Figure 1:**
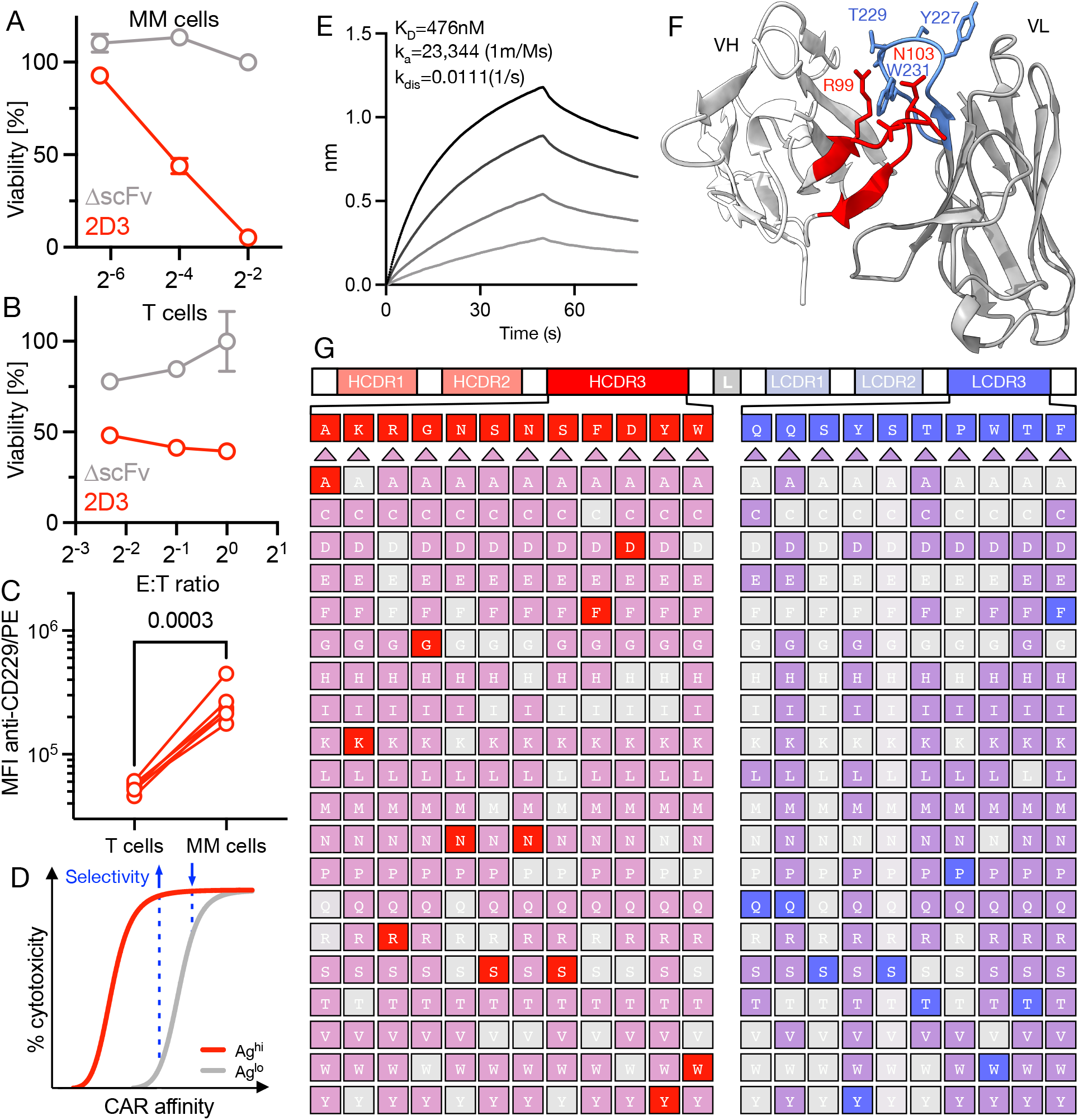
Generation of 2D3-based CDR3 variant library for the generation of low affinity CD229 antibodies with increased selectivity. **(A)** Killing of CD229^pos^ MM cell line U266B1 expressing luciferase by CD229 CAR T cells (2D3) or T cells expressing a CAR without a binding domain (ΔscFv) as determined by luminescence assay. Data indicate mean ± S.D. from technical replicates (*N*=3). **(B)** Killing of healthy human T cells by CD229 CAR T cells (2D3) as determined by flow cytometry cytotoxicity assay. Data indicate mean ± S.D. from technical replicates (*N*=3). **(C)** Surface expression of CD229 on MM and T cells from patients with relapsed/refractory MM (*N*=6). Expression was determined after staining with an anti-CD229 antibody (clone: HLy9.1.25) on a CytoFLEX LX flow cytometer (Beckman Coulter). **(D)** Schema of relationship between CAR affinity and targeting of cells expressing high antigen levels (Ag^hi^) and low antigen levels (Ag^lo^). **(E)** Sensorgram of 2D3 binding to CD229. Equilibrium and rate constants of the 2D3 scFv were determined by biolayer interferometry (BLI). Biotinylated 2D3 was immobilized on a streptavidin biosensor and the recombinant extracellular domain of CD229 was added in the following concentrations: 2μM, 1μM, 0.5μM, 0.25μM. Sensorgram indicates binding curves for descending CD229 concentrations. Plot shows a representative result of three independent experiments. **(F)** Structure of the 2D3 scFv as predicted by AlphaFold2 with the GS-linker omitted. CDR3 loops of the variable heavy (red) and the variable light (blue) chains as well as exposed residues are highlighted. **(G)** Schema of single amino acid substitution 2D3 CDR3 variant library indicating represented residues. Missing variants (grey) and wild-type residues (red/blue) are highlighted.

### Single amino acid substitutions result in substantially reduced CD229 binding

Identification of antibodies with reduced affinity represents a relatively uncommon objective in antibody discovery and poses unique challenges when developing appropriate screening approaches. Most common primary antibody screening assays, such as standard solid-phase binding assays using large sets of non-purified antibodies, are unable to differentiate between expression and affinity. As expected, this was also the case for 2D3, which showed a clear dependence of CD229 binding to antibody concentration, especially in concentration ranges commonly observed in standard high-throughput expression cultures (Fig. 2A). When screening for high-affinity antibodies this may be acceptable as the highest assay signals are likely the result of a combination of high affinity and high expression, both representing desirable properties. In the case of low affinity antibody screening, however, the potential conflation of low expressing/high affinity antibodies with high expressing/low affinity antibodies would render such data relatively meaningless. We therefore developed a high-throughput scFv quantification assay relying on the binding of Protein L to the κ light chain in 2D3 (Fig. 2B) and determined total scFv yields of all generated variants (Fig. 2C). As expected, substitutions to cysteine and proline generally resulted in poor antibody expression, while alanine and threonine appeared to be tolerated well in various positions (Fig. 2C). In addition, we found that some positions were able to accommodate almost any amino acid. This included some unexpected positions, such as asparagine in position H5, an exposed residue in the center of the groove formed by both CDR3 loops, as well as tryptophan in H12 and threonine in L9 (Fig. 2C). Importantly, several substitutions resulted in substantially improved expression, in line with established approaches to improve antibody stability via mutation of non-contact residues ^23, 24^. Following normalization of antibody concentrations, we next determined binding to recombinant CD229 in a standard solid-phase binding assay (Fig. 2D). While the majority of mutations did not reduce 2D3 variant binding to CD229, and some, as expected, increased binding, the comprehensive mutagenesis approach taken enabled the identification of a large set of binders showing various levels of reduced binding. While mutation of the outer residues in both CDR3s appeared to most drastically reduce binding, mutation of any position affected binding to some extent following individual amino acid substitutions. Importantly, alanine scanning^25^, one of the most widely used approaches to reduce protein-protein binding would not have resulted in a similarly comprehensive set of variants as alanine substitutions in many cases did not alter binding compared to parental 2D3 when other substitutions substantially affected binding. In addition, alanine substitutions never represented the variants with the lowest binding signal in any position.

**Figure 2:**
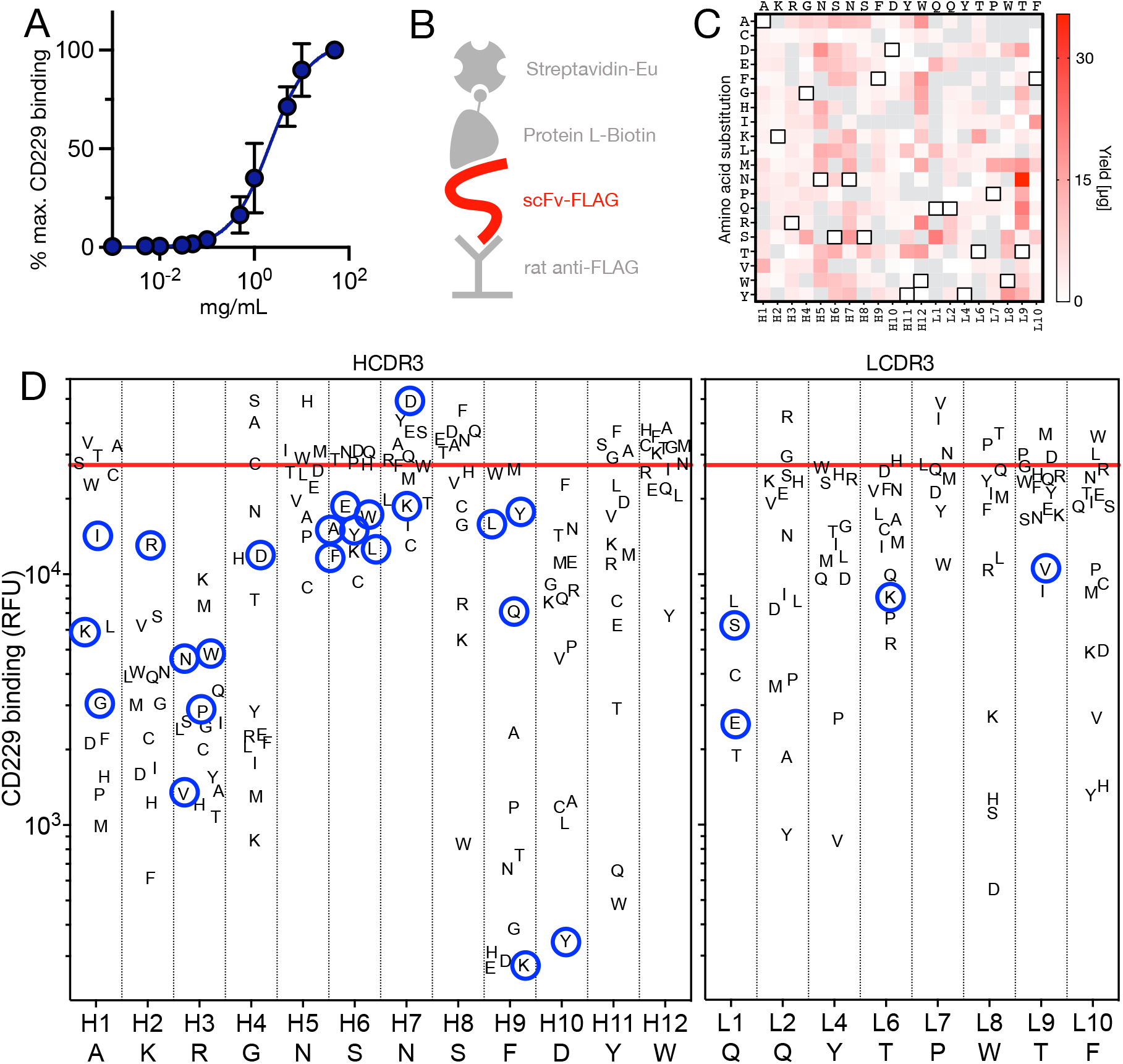
Single amino acid substitutions result in substantially reduced CD229 binding. **(A)** Concentration-dependence of 2D3 scFv binding to recombinant CD229 as determined by solid-phase time-resolved fluorescence assay. Data indicate mean ± S.D. from technical replicates (*N*=3). **(B)** Schema of solid-phase assay for the determination of FLAG-tagged scFv concentrations using rat IgG2a 11 anti-FLAG antibody, biotinylated Protein L, and streptavidin-Eu. **(C)** Total yield of 2D3 variants from autoinduction cultures as determined by assay illustrated in 2B. White squares with black outlines indicate amino acids in wildtype 2D3. Plot shows representative result of two independent experiments. **(D)** Binding of all expressed 2D3 variants at 2ng/μl to recombinant CD229 as determined by solid-phase time-resolved fluorescence assay. Red line indicates wildtype 2D3 binding to CD229. Blue circles indicate variants selected for downstream assays based on amino acid position and binding signal. Plot shows representative result of two independent experiments.

Taken together, our data indicate that comprehensive single amino acid substitution may be preferable to conventional mutagenesis strategies and is able to generate large sets of antibodies showing reduced antigen binding even in already relatively low affinity binders. Based on scFv expression and CD229 binding data, we next selected 26 binders for downstream analyses.

### Affinity tuning approach results in predominantly off-rate driven affinity reductions

To determine if single amino acid substitution mutagenesis in fact resulted in relevantly altered affinities and rate constants, we next purified (Suppl. Fig. 2) *in vivo* biotinylated 2D3 variant scFvs (Fig. 3A). We then subjected these antibodies to biolayer interferometry (BLI) using streptavidin biosensors and recombinant CD229 (Fig. 3B), allowing the use of relatively high analyte concentrations due to non-destructive sampling^26^, which facilitated characterization even of very weak binders (Fig. 3C). Affinities of 2D3 variants with single amino acid substitutions ranged from 175nM to >10,000nM (Table 1) and we observed a very close correlation between antibody affinity (K_D_) and TRF binding (Suppl. Fig. 3). Differences in affinity were generally driven by faster off-rates (Table 1), which is likely related to the use of a solid phase binding assay for primary variant screening, which may have biased clone selection towards variants with faster off-rates. In contrast to this finding, we observed a dramatic reduction in the on-rate of variants in which arginine in H3 was replaced (Fig. 3C, Table 1). A noted exception to this finding is RH3V, which in fact showed a faster on-rate but also a much faster off-rate (Fig. 3D). These data suggest a key role of arginine in the orchestration of the 2D3 epitope and among other possibilities might point to a model of 2D3 binding to CD229 in which RH3 binding facilitates interactions by other residues.

**Figure 3:**
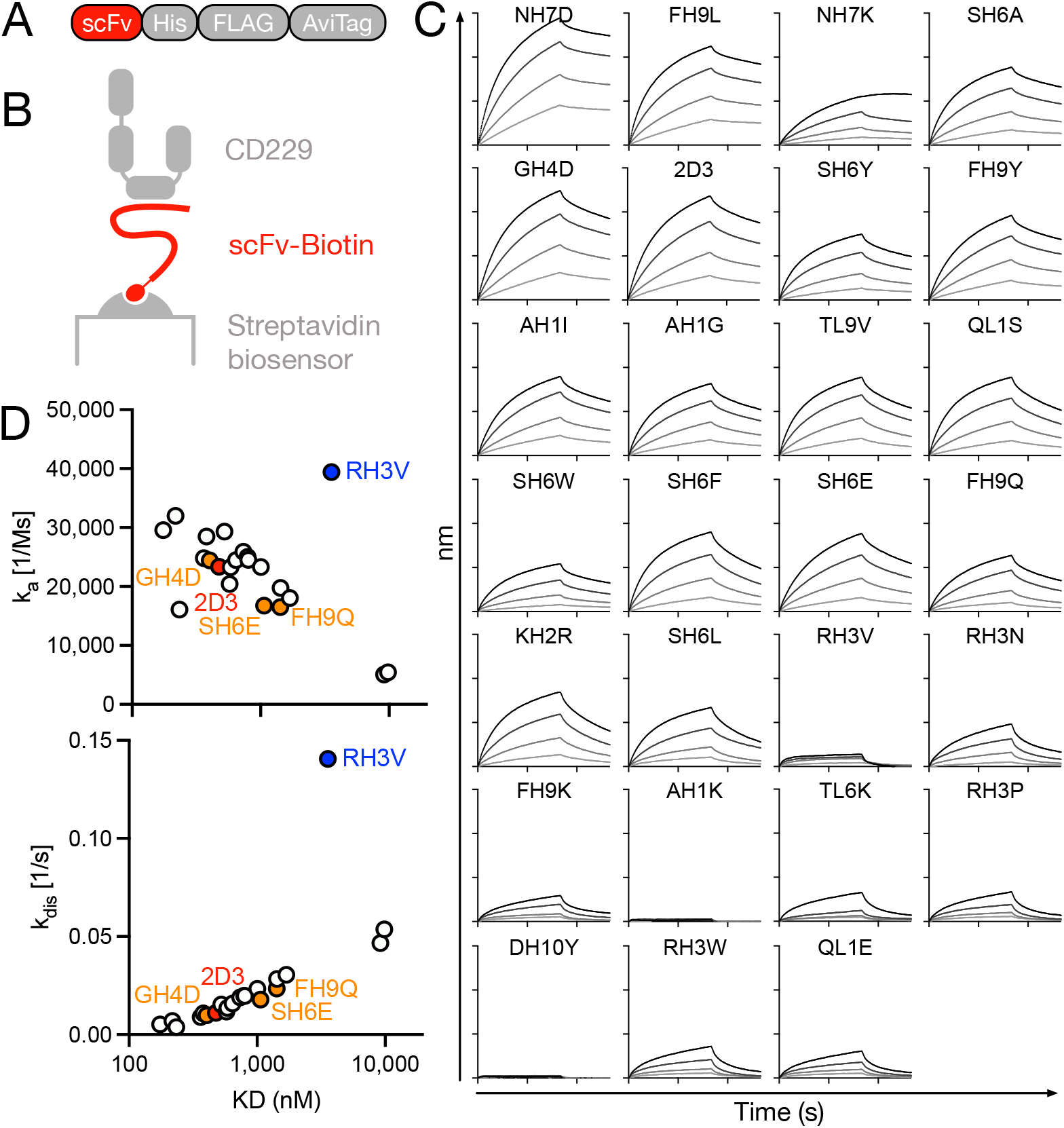
Affinity tuning approach results in predominantly off-rate-driven affinity reductions. **(A)** Schema of construct used for production of biotinylated 2D3 scFv variants including C-terminal Avitag to facilitate *in vivo* biotinylation. **(B)** Schema of biolayer interferometry (BLI) setup used for kinetic characterization of CD229 binding. Biotinylated 2D3 variants were immobilized on streptavidin biosensors and the recombinant extracellular domain of CD229 was added in the following concentrations: 2μM, 1μM, 0.5μM, 0.25μM. **(C)** Sensorgrams of 2D3 variants were determined using an Octet K2 (Sartorius). Plots show representative result of two independent experiments. **(D)** Correlation between rate and equilibrium constants of 2D3 variant scFvs as determined by BLI.

**Table 1:** Single amino acid substitution results in broad range of affinities. Equilibrium and rate constants of 2D3 variants were determined using an Octet K2 (Sartorius). Data are representative of two independent experiments. Parental 2D3 is shown in red and variants with increased selectivity are shown in orange.

| Clone | KD<br>(nM) | ka (1/M.s) | kdis (1/s) |
| --- | --- | --- | --- |
| NH7D | 175 | 29590 | 0.0052 |
| FH9L | 218 | 32007 | 0.0070 |
| NH7K | 234 | 16084 | 0.0038 |
| SH6A | 380 | 28502 | 0.0108 |
| GH4D | 402 | 24421 | 0.0098 |
| 2D3 | 476 | 23344 | 0.0111 |
| SH6Y | 524 | 29323 | 0.0154 |
| FH9Y | 575 | 20420 | 0.0117 |
| AH1I | 585 | 23268 | 0.0136 |
| AH1G | 644 | 24487 | 0.0158 |
| TL9V | 736 | 25885 | 0.0190 |
| QL1S | 791 | 25039 | 0.0198 |
| SH6W | 803 | 24513 | 0.0197 |
| SH6F | 1,006 | 23307 | 0.0234 |
| SH6E | 1,065 | 16737 | 0.0178 |
| FH9Q | 1,425 | 16507 | 0.0235 |
| KH2R | 1,431 | 19747 | 0.0283 |
| SH6L | 1,689 | 18046 | 0.0305 |
| RH3V | 3,565 | 39427 | 0.1406 |
| RH3N | 9,157 | 5088 | 0.0466 |
| FH9K | 9,835 | 5452 | 0.0536 |
| AH1K | >10,000 | 50315 | 0.9477 |
| TL6K | >10,000 | 1337 | 0.0635 |
| RH3P | >10,000 | 755 | 0.0679 |
| DH10Y | >10,000 | 8164 | 0.8991 |
| RH3W | >10,000 | 595 | 0.0703 |
| QL1E | >10,000 | 558 | 0.0688 |

Single amino acid substitution together with BLI not only resulted in the identification of a set of clones with a wide range of mostly off-rate driven differences in affinity but also provided data regarding the mode of 2D3-CD229 binding, which may aide further lead optimization.

### CD229 CAR T cells based on variant antibodies can be manufactured, show efficient CAR surface expression, and allow identification of clones with increased selectivity

Optimal CAR affinity remains an active area of research but likely depends on various parameters, such as need for selectivity, epitope, as well as antigen- and CAR-density. Ideally, CAR affinity for a given target antigen will be chosen empirically, thus requiring a sufficiently large set of binders with different affinities available for CAR construction. We therefore converted all 26 antibodies into CAR constructs (Fig. 4A) and produced primary human CAR T cells using a standard manufacturing process (Fig. 4B). It has previously been shown that even parental 2D3-based CAR T cells can be manufactured without the CAR T cells targeting each other because CD229 is downregulated upon CD3/CD28-bead activation at the start of manufacturing^21^. We therefore first determined the viability and total CAR T cell yields on day 7 of manufacturing and found that yields varied substantially between constructs (Fig. 4C), possibly indicating increased tonic signaling ^27, 28^. Considering the substantial differences in scFv expression levels between variants (Fig. 2C), we next determined CAR surface expression of wildtype 2D3 and all variant CAR constructs, as well as T cells expressing a CAR without a binding domain (ΔscFv). While all variant constructs expressed similar levels of the linked GFP reporter, two constructs, FH10K and AH1K, showed relatively low CAR surface expression and two other constructs, DH10Y and RH3N, did not show any CAR surface expression at all (Fig. 4D). One construct, TL9V, showed a bimodal distribution, potentially indicating recombination during retroviral packaging. We next determined whether mutagenesis had resulted in altered anti-tumor activity by determining killing of MM cells by all variant CAR T cells at multiple effector-target ratios. We found that while several constructs indeed showed reduced tumor cell killing, likely due to the substantially reduced affinity of those variants, several constructs, including SH6A, GH4D, FH9Y, AH1I, AH1G, and FH9Q showed either equal or enhanced anti-tumor activity (Fig. 4E, Suppl. Fig. 4). Next, we set out to determine whether reducing CAR affinity resulted in increased selectivity by measuring killing of purified T cells (Suppl. Fig. 5). At an effector-target ratio of 0.5:1, we found that, as expected, numerous constructs that had shown reduced cytotoxic activity against tumor cells, now also showed reduced killing of T cells (Fig. 4F). However, among the variants showing comparable or increased tumor cell killing, we were able to identify one construct, FH9Q, which showed dramatically reduced killing of T cells (Fig. 4F).

**Figure 4:**
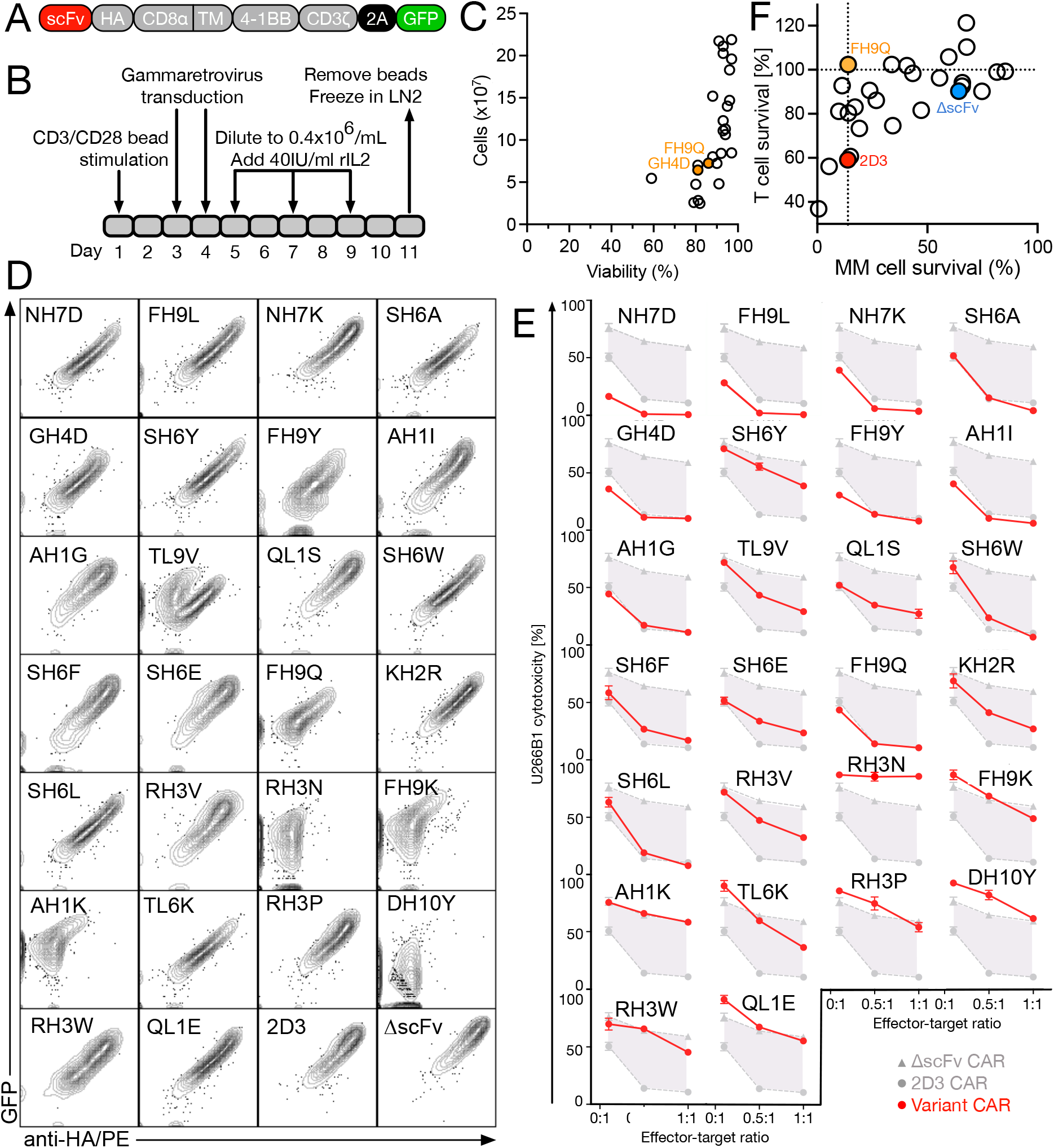
Multiple HCDR3 variants maintain anti-tumor activity but exhibit minimal T cell killing. **(A)** Schema of 4-1BB-based second-generation CAR construct with GFP reporter. **(B)** Schema of gammaretrovirus-based CAR T cell production process. **(C)** Correlation between CAR T cell yield and viability of 26 2D3 variant CARs. **(D)** Surface expression of 2D3 variant CARs and GFP reporter expression as determined by anti-HA staining using flow cytometry. **(E)** Cytotoxic activity of 2D3 variant CAR T cells against MM cell line U266B1 expressing luciferase at different effector-target ratios using a luminescence-based cytotoxicity assay. Data indicate mean ± S.D. from technical replicates (*N*=3). **(F)** Correlation of cytotoxic activity of 2D3 variant CAR T cells against MM cells and T cells overnight at an effector-target ratio of 0.5:1. Data indicate mean killing (fold of wildtype 2D3) ± S.D. from technical replicates (*N*=3). Candidates were selected for downstream assays based on increased selectivity and anti-tumor activity (orange).

### FH9Q CAR T cells maintain anti-tumor activity in vitro and in vivo, do not target healthy T cells, and evidence reduced trogocytosis

Although we had hypothesized that a single amino acid substitution would be unlikely to substantially alter the specificity of the binding domain as a whole, we next determined whether FH9Q-based CAR T cells would still specifically target cells expressing CD229. We indeed found that FH9Q CAR T cells did not kill CD229^neg^ K562 cells but showed substantial killing of K562 cells engineered to express CD229 (Fig. 5A). When scaling up FH9Q CAR T cell production, we observed that FH9Q CAR T cells, similar to many of the other variants (Fig. 4C) showed reduced expansion compared to ΔscFv CAR T cells *in vitro* (Fig. 5B) and *in vivo* (Fig. 5C), which had not been the case for parental 2D3-based CAR T cells. These results are consistent with a previous report describing reduced *in vitro* and *in vivo* expansion of CAR T cells evidencing tonic signaling ^27^ – the spontaneous activation of T cells by CARs in the absence of antigen, likely as a consequence of aggregation mediated by the CAR binding domain. Tonic signaling has been shown to lead to the rapid depletion of CJUN, a component of the T cell activating AP-1 transcription factor. and overexpression of CJUN was shown to efficiently rescue function and expansion of CAR T cells evidencing tonic signaling ^28^. We therefore generated an FH9Q CAR construct to simultaneously overexpress CJUN (Fig. 5D) and found that CJUN overexpression efficiently restored FH9Q CAR T cell expansion *in vitro* (Fig. 5B) and *in vivo* (Fig. 5C). Because we suspected that the FH9Q CAR may be prone to aggregation, we next analyzed its expression on the surface of T cells, because it had previously been shown that antibodies with higher levels of aggregation exhibit reduced surface expression ^29^. We indeed observed substantially reduced FH9Q CAR expression compared to 2D3, suggesting increased aggregation of FH9Q-based CARs (Suppl. Fig. 7). We further speculated that this difference in CAR surface expression may have contributed to the cells’ increased selectivity. To determine the influence of surface expression levels on selectivity, we next sorted 2D3, ΔscFv, and FH9Q CAR T cells to normalize for CAR surface expression (Suppl. Fig. 7). Comparing sorted and unsorted 2D3 CAR T cells, we did, however, not observe differences in their ability to kill MM cells (Suppl. Fig. 8). We next compared the ability of FH9Q and 2D3 CAR T cells sorted for comparable CAR surface expression to kill MM cells and found that killing by these cells remained identical as well (Fig. 5E). Importantly, in a direct co-culture, we observed significantly reduced killing of healthy T cells by FH9Q CAR T cells compared to 2D3 CAR T cells even following normalization of CAR surface expression (Fig. 5F). These data suggest, that altered surface expression levels did not meaningfully contribute to the increased selectivity of FH9Q CAR T cells, which is therefore likely the result of their reduced affinity.

**Figure 5:**
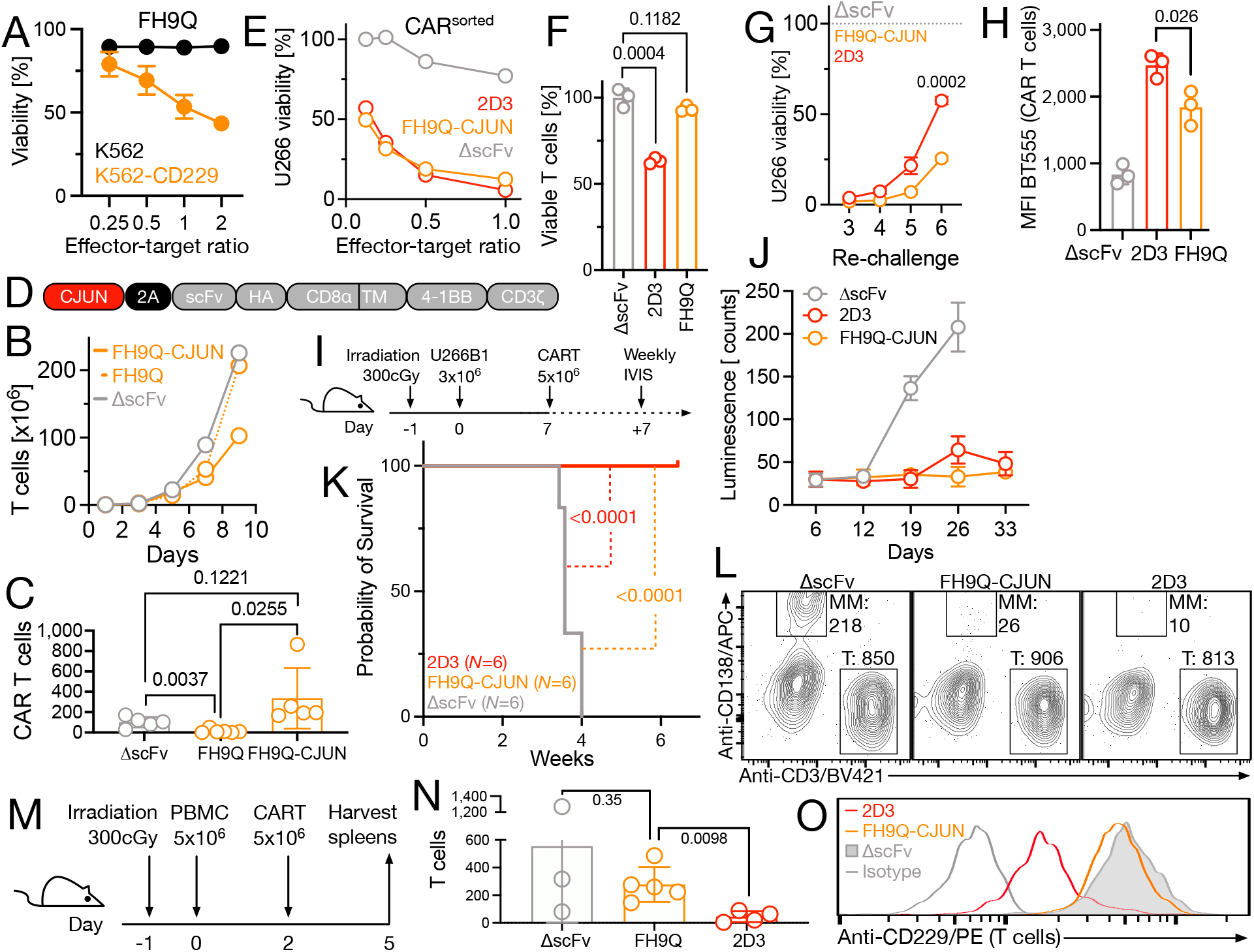
Low affinity variants exhibit minimal T cell killing and reduced trogocytosis, while maintaining target specificity and anti-MM activity. **(A)** Killing of parental CD229-negative K562-Luc cells and K562-Luc cells transduced with a CD229 expression construct. Target cell killing was determined by luciferase-based cytotoxicity assay. Data indicate mean ± S.D. from technical replicates (*N*=3). **(B)** Expansion of CD229 CAR T cells during manufacturing as determined by cell counting. Data are representative of 2 independent experiments. **(C)** NRG mice bearing U266B1 tumors were injected with T cells expressing FH9Q CAR T cells with or without CJUN. Animals were euthanized between days 7 and 9 and CAR T cell numbers determined by flow cytometry. Data indicate mean ± S.D. from 5 animals per group. Statistical differences between conditions were determined by two-sided Student’s t test. **(D)** Retroviral construct used for the simultaneous expression of CARs and CJUN. **(E)** Killing of U266B1-Luc cells by HA^int^ sorted CD229 CAR T cells after an overnight co-culture as determined by luminescence cytotoxicity assay. Data indicate mean ± S.D. from technical replicates (*N*=3). **(F)** Killing of healthy autologous T cells by CD229 CAR T cells in an overnight in vitro co-culture at an effector-target ratio of 1:1. Data indicate mean ± S.D. from technical replicates (*N*=3). Statistical differences between conditions were determined by two-sided Student’s t test. **(G)** Repeated killing of U266B1-Luc cells in an in vitro overnight co-culture assay by CD229 CAR T cells after daily rechallenge with tumor cells. Data indicate mean ± S.D. from technical replicates (*N*=3). **(H)** Membrane transfer from U266B1 cells to variant CAR T cells after 4h co-culture at an effector-target ratio of 4:1 as determined following Biotracker 555 staining of U266B1 cells using flow cytometry. Data indicate mean ± S.D. from technical replicates (*N*=3). Statistical differences between conditions were determined by two-sided Student’s t test. **(I)** Schema of in vivo experiment to determine the efficacy of low affinity CD229 CAR T cells. **(J)** Bioluminescence of mice was determined using an in vivo imaging system (IVIS). Data indicate mean ± S.D. from 6 animals per group. **(K)** Cumulative survival of NSG mice injected i.v. with 3×10^6^ U266B1 cells on day 0 and 5×10^6^ CD229 CAR T cells on day 7. Statistical significance was determined by log-rank test. **(L)** Overnight cytotoxicity assay to determine relative targeting of MM and healthy T cells by CD229 CAR T cells using flow cytometry. 5×10^4 T cells and 5×10^4 U266B1 cells were cocultured with 1×10^5 CD229 CAR T cells and relative killing was determined by flow cytometry. Numbers indicate total cell numbers within the respective gates normalized using counting beads. **(M)** Schema of short-term *in vivo* experiment to determine targeting of healthy T cells by CD229 CAR T cells. **(N)** Numbers of CD3^pos^ HA/CAR^neg^ healthy T cells as determined by flow cytometry per 50,000 events in spleens from mice injected with 5×10^6^ purified PBMCs and subsequently treated with CD229 CAR T cells. Data indicate mean ± S.D. from 3-5 individual animals. Statistical differences between conditions were determined by two-sided Student’s t test. **(O)** Surface expression of CD229 on healthy T cells after co-culture with indicated CAR T cells for 8h at an effector target ratio of 5:1 as determined by flow cytometry.

We next asked the question whether the substantial reduction in CAR affinity would result in incomplete tumor cell killing or suboptimal CAR T cell stimulation leading to reduced long-term disease control by FH9Q CAR T cells. We therefore first performed an *in vitro* re-challenge assay and observed that FH9Q CAR T cells in fact showed significantly increased long-term disease control compared to 2D3 CAR T cells (Fig. 5G). This finding may be related to the overexpression of CJUN together with FH9Q or more physiological T cell stimulation by low affinity CAR constructs^13^, but could also be the result of reduced trogocytosis by low affinity CAR T cells, which we have previously shown to enhance CAR T cell persistence ^30^. Trogocytosis is the stripping of target antigen together with target cell membrane and their incorporation into the CAR T cell membrane, resulting in antigen negative tumor cells and antigen-positive CAR T cells ^30, 31^, potentially leading to fratricide, the killing of CAR T cells by other CAR T cells. We therefore determined the level of membrane transferred from MM cells to CAR T cells in a short-term co-culture. We found that FH9Q CAR T cells had indeed transferred significantly less tumor membrane than 2D3 CAR T cells (Fig. 5H), which may in turn have contributed to improved CAR T cell persistence and long-term disease control. In a mouse model of human MM (Fig. 5I), we also observed that both 2D3 and FH9Q CAR T cells significantly reduced tumor burden (Fig. 5J) and prolonged survival (Fig. 5K), indicating that FH9Q CAR T cells show long-term anti-tumor activity comparable to 2D3 CAR T cells. In addition, we observed comparable levels of major effector cytokines between 2D3 and FH9Q CAR T cells when co-cultured with MM cells, indicating potent T cell activation by FH9Q CAR T cells (Suppl. Fig. 9).

As killing of healthy T cells represents the main liability of CD229 CAR T cells, we next explored the apparent lack of targeting of healthy T cells by FH9Q CAR T cells. Interestingly, while we had observed significant killing of healthy T cells in a direct-co-culture by 2D3 CAR T cells (Fig. 5F), we only found minor T cell killing in a mixed co-culture containing both MM and healthy T cells as targets, suggesting that 2D3 CAR T cells already provide a degree of selectivity in the presence of tumor cells (Fig. 5L). However, in the absence of MM cells, we not only observed substantial killing *in vitro* (Fig. 5F) but also in an *in vivo* cytotoxicity assay (Fig. 5M/N). Importantly, FH9Q CAR T cells did not show killing of healthy T cells in any of our *in vitro* and *in vivo* assays. Previously, we had observed that treatment with 2D3 CAR T cells results in a population of CD229^low^ healthy T cells *in vitro* and *in vivo* ^21^. Analyzing CD229 expression on healthy T cells following co-culture with 2D3 or FH9Q CAR T cells, we again observed emergence of a CD229^low^ T cell population when treated with 2D3 CAR T cells. However, we did not observe selection of a CD229^low^ population when treated with FH9Q CAR T cells, further substantiating the lack of targeting of healthy T cells (Fig. 5O). Overall, our data demonstrate that FH9Q CAR T cells maintain the anti-tumor activity of 2D3 CAR T cells *in vitro* and *in vivo* but lack their cytotoxic activity against healthy T cells.

## DISCUSSION

CAR T cells have revolutionized cancer immunotherapy, however, the development of safer and more effective CAR T cell approaches will be critical for their more widespread adoption and increased patient benefit. CAR affinity has recently come into focus due to increased tumor selectivity and improved CAR T cell function resulting from the use of low affinity CAR constructs^13–16^. In addition, affinity tuning of CAR constructs opens the door to the targeting of new antigens that would otherwise be excluded due to their expected toxicity profile. Here, we provide a systematic approach for the development of minimally modified low affinity antibody variants based on established and extensively tested antibodies that does not require detailed structural information including the antibody’s exact epitope. We demonstrate that the variants produced using this approach indeed show significantly increased selectivity and improved CAR T cell function, while maintaining the original epitope and specificity. The majority of variants generated using our approach show a predominantly off-rate based reduction in affinity, likely as a consequence of using a solid phase binding assay for primary variant screening. While data regarding the relative contribution of on-rate and off-rate on CAR T cell signaling remain scarce, prior approaches to modulate CAR affinity have predominantly focused on changes in off-rates^32–34^. This includes one of the most advanced approaches, which is already demonstrating the clinical benefits of the use of low affinity CAR constructs targeting CD19 ^13, 33^.

We show that a potential liability of our approach is the potential for tonic signaling in variant CAR T cells despite only minor changes to the binding domain. However, as shown here, at least in some cases it may be relatively simple to overcome this issue by rendering the CAR T cells resistant to some of the downstream effects of tonic signalling by overexpressing CJUN. An alternative strategy to address the problem of tonic signaling and to improve CAR T cell function in general is an elegant novel approach using protease-sensitive CAR constructs ^35^. In this system, small molecule-mediated protease inhibition allows tight control over CAR surface expression levels, preventing early exhaustion resulting from tonic signaling, maintenance of a stem-like phenotype, and dramatically increased anti-tumor activity even when using aggregation-prone CAR constructs.

Another drawback of our approach could be that parental antibodies with very high affinities may require multiple iterations of mutagenesis to achieve a sufficient reduction in affinity and that for some parental antibodies it may not be possible to generate variants within the required affinity window depending on their binding mode. However, we show that our combinatorial approach is likely to produce a wide range of affinities and represents a robust workflow for the development of safer and more functional CAR T cells.

## METHODS

### Cell lines and primary human cells

U266B1, K562, and Phoenix-Ampho cells were purchased from ATCC and cultured according to ATCC instructions. Lenti-X 293T cells were purchased from Takara and cultured according to the manufacturer’s instructions. Cell lines were authenticated by their respective supplier. Healthy donor buffy coats were obtained from the Blood Centers of America and the New York Blood Center and peripheral blood mononuclear cells were isolated from buffy coats by density gradient using FicollPaque (GE) as previously described ^21^.

### Flow cytometry expression analysis

Flow cytometry staining and analyses were performed as previously described ^21^. CD229 surface expression was determined using a mouse monoclonal anti-CD229 antibody (clone: HLy9.1.25). Other antibodies used for flow cytometry analyses are listed in Supplementary Table 1. Commercially available antibodies were used at dilutions recommended by the respective manufacturer. For the analysis of CD229 expression on tumor cells and T cells from MM patients, data was acquired on a CytoFLEX LX (Beckman Coulter) and analyzed using Kaluza 2.1 (BC). All other flow cytometry data was acquired on an LSR Fortessa or LSR II flow cytometer (BD) and analyzed using FlowJo 10 (BD).

### Single amino acid substitution library production

Parental 2D3 was cloned into pSANG10 ^36^ and the single amino acid substitution library produced by was generated using high-throughput gene synthesis by Twist Bioscience. Individual mutations were confirmed by Sanger sequencing. To produce single-site biotinylated scFvs, a C-terminal AviTag was added to the pSANG10 expression constructs (pSANG10-Avi) and scFvs transformed into BL21[DE3] cells (Lucigen) containing the pBirAcm plasmid (Avidity).

### scFv expression and purification

Monoclonal scFvs were expressed overnight in 25 mL MagicMedia E. coli autoexpression medium (Thermo-Fisher). Periplasmic extracts were generated from autoinduction cultures using standard procedures. For some experiments, scFvs were purified by metal affinity chromatography using NiNTA resin (Thermo). Concentrations of purified antibodies were determined by bicinchinonic assay (Thermo). pSANG10-Avi clones were expressed in the same way in the presence of 50μM free D-biotin (Sigma).

### IFN gamma ELISA

Supernatants were harvested from 96 well overnight co-cultures and immediately frozen at −80°C. IFNγ concentrations were determined via standard curve using a commercial ELISA kit according to the manufacturer’s instructions (Biolegend). Absorbance was measured on a multi-mode plate reader (Tecan).

### CAR T cell production

Selected scFvs were cloned into a previously described second generation 4-1BB-based CAR construct^21^ in the gammaretroviral SFG backbone. Amphotropic gammaretrovirus was generated by transfection of Phoenix-Ampho cells (ATCC # CRL-3213) using the SFG-based transfer plasmids using Lipofectamine 2000 according to the manufacturer’s instructions. Virus-containing supernatants were concentrated with Retro-X concentrator (Takara). PBMCs were stimulated for 2 days with CD3/CD28 T cell activation beads (Thermo # 11131D) in the presence of 40IU/mL IL2 (R&D Systems # 202-IL-010) in AIM V (Thermo) supplemented with 5% human serum (Sigma #H3667) and incubated at 37°C/5% CO_2_. Bead-stimulated cells were transferred to Retronectin-coated (Takara) virus-containing plates and incubated overnight. Transduction was repeated the next day before counting and diluting cells to 0.4×10^6^ cells/ml. After the second transduction cells were grown for an additional 7 days before removing beads using a DynaMag-15 magnet (Thermo). IL-2 was replenished every 2 days to 40IU/mL. Cells were frozen in 90% FCS/10% DMSO and stored in liquid nitrogen.

### Trogocytosis assay

CAR T cells were cocultured with target cells at the specified effector:target ratio. Target cells were first labelled with BioTracker 555 (Sigma # SCT107) according to the manufacturer’s instructions. After the specified amount of time, cells were stained with antibodies and 500ng/mL 4′,6-diamidine-2′-phenylindole dihydrochloride (DAPI, Invitrogen # D1306). Samples were analyzed on an LSR II flow cytometer (BD).

### Flow cytometry-based cytotoxicity assay

A flow cytometry-based cytotoxicity assay was used to determine CAR T cell cytotoxicity against healthy T cells from the same healthy donor as well as primary MM cells. T cells were collected using negative selection (Stemcell Technologies EasySep Human T Cell Isolation Kit) from autologous healthy donor PBMCs. MM cells and T cells were stained with Cell Trace Far Red dye (CTD, Invitrogen) according to the manufacturer’s instructions. 5×10^4^ target cells were co-cultured with different amounts of CAR T cells overnight in a round bottom 96 well plate at 37°C 5%CO2. Following coculture, Accucheck counting beads (Life Technologies) and 500ng/mL DAPI were added to the cells. DAPI^-^CTD^+^ T Cells were immediately quantified on an LSR II flow cytometer (BD).

### Luciferase-based cytotoxicity assay

To determine the cytotoxicity of variant CD229 CAR T cells against the multiple myeloma cell line U266B1 and chronic lymphocytic leukemia cell line K562, cell lines were transduced with pHIV-Luc-ZsGreen lentivirus and sorted on a FACSaria flow cytometer (BD) for GFP expression. CD229-negative K562 cells were also transduced with a CD229 expression construct as previously described ^21^. As with the flow cytometry-based cytotoxicity assay, 5×10^4^ target cells were seeded in each well of a round bottom 96 well plate. Various ratios of CAR T cells were co-cultured with target cells overnight at 37°C 5% CO_2_. After the co culture, cells were suspended by gentle pipetting and 100uL were moved to a 96 well black flat bottom plate. 150 μg/ml D-luciferin (Gold Biotechnology Cat# LUCNA-2G) was added to the cells and incubated for 5 mins at 37°C. Luminescence was determined on a multi-mode plate reader (Tecan Spark). For the re-challenge assay, luminescence was determined daily before adding 5×10^4^ to each well.

### Time-resolved fluorescence assays

Two different time-resolved fluorescence (TRF) assays were used. To determine the concentration of scFv in the PPEs, 250ng of rat-anti-FLAG (clone: L5, Biolegend) in 50uL PBS was immobilized on a black 96-well plate (Greiner Bio-One) overnight at 4°C. Plates were washed using an automated plate washer (Tecan HydroFlex) twice with PBS containing 0.1% Tween-20, and twice with PBS, in between each incubation. After immobilization, all other incubations were performed at room temperature at 400rpm. Following immobilization, plates were blocked in 3% non-fat milk in PBS (M-PBS). Then PPEs or purified 2D3 were added to plates in 3% M-PBS and incubated for 1h. Next, plates were incubated with 250ng biotinylated protein L (Thermo Scientific) in 3% M-PBS for 1h. Finally, plates were incubated with streptavidin-Europium (PerkinElmer) in PBS for 30min. After a final wash, plates were incubated with DELFIA Enhancement solution (PerkinElmer) for 10min. Time-resolved fluorescence was determined on a multi-mode plate reader (Tecan Spark). A purified parental 2D3 standard was used to calculate the scFv concentration in each PPE.

To determine the relative binding of variant scFvs to CD229, 5 μg/ml recombinant human CD229 (R&D Systems) was immobilized on a black 96-well plate (Greiner Bio-One) overnight at 4°C. After immobilization, plates were washed between each step and incubated at room temperature at 400rpm as described above. Following immobilization, plates were blocked in 3% M-PBS. Then, plates were incubated with 100ng of each scFv in 3% M-PBS. Next, plates were incubated with anti-FLAG M2 (Sigma-Aldrich) in 3% M-PBS. Finally, plates were incubated with anti-mouse IgG-Europium antibody (PerkinElmer) for 1 hour. After a final wash, plates were incubated with DELFIA Enhancement solution (PerkinElmer) for 10min. Time-resolved fluorescence was determined on a multi-mode plate reader (Tecan Spark).

### Biolayer Interferometry (BLI)

Streptavidin (SA) biosensensors (FortéBio) were hydrated in 1x Octet® Kinetics Buffer (Sartorius) for at least 10min. SA biosensors were loaded using the Octet K2 (Sartorius). A baseline in Octet Kinetics Buffer was collected for 1 min. Then the sensors were loaded with variant scFvs using a threshold of 2nm and subsequently blocked in Biocytin. Once loaded, sensors were placed back into the sensor tray and kept hydrated in kinetics buffer. Once sensors were loaded, a kinetic run was performed. SA biosensors loaded with 2nm of biotinylated scFv and blocked with biocytin went through a 60s baseline read in kinetics buffer, 50s association in 2, 1, 0.5, and 0.25µM recombinant human CD229 (R&D Systems) in kinetics buffer, and finally 60s dissociation in kinetics buffer. BLI was run at 30°C and 10,000rpm. Data was analyzed using Octet^®^ K2 System Data Analysis 9.0 software.

### 2D3 structure prediction

The structure of the wildtype 2D3 scFv was generated using AlphaFold2 using default parameters ^37^. Structures were visualized using UCSF ChimeraX.

### Multiple myeloma xenograft mouse model

Six- to 8-week-old male NOD.Cg-Rag1^tm1Mom^ Il2rg^tm1Wjl^/SzJ (NRG, The Jackson Laboratory (Cat#005557)) mice were sublethally irradiated with 300cGy (Rad-Source RS-2000) and injected on the next day via the lateral tail vein with the indicated numbers of U-266 stably expressing luciferase. On day 7 after tumor cell injection, the indicated numbers of CD229 CAR T cells or CAR T cells lacking a binding domain (ΔscFv) were injected into the tail vein. Animals were weighed twice weekly and monitored for signs of distress in accordance with institutional regulations. For in vivo imagining, mice received an intraperitoneal injection of 3.3mg D-luciferin. Photographic and luminescent images were acquired starting 10min after the D-luciferin injection, both in prone and supine position using a IVIS imaging system. Myeloma progression was monitored every 7 days until the study endpoint. Average radiance (p/s/cm²/sr) was quantified for individual animals using Living Image software (PerkinElmer).

### In vivo cytotoxicity assay using human PBMCs

Eight-week-old male NRG mice were sublethally irradiated with 300cGy and on the following day injected with 5×10^6^ PBMCs isolated from healthy donors. On day 2 after PBMC injection, mice were injected with 5×10^6^ CD229 CAR T cells via tail vein. On day 5 after PBMC injection, animals were sacrificed and spleens were collected for flow cytometry analysis. After a 5min incubation in red blood cell lysis buffer (Biolegend), cells were washed twice in PBS, incubated with human and mouse FcR blocking reagents (Miltenyi Biotec) for 15min, and then stained with population-specific antibodies (Supplementary Table 1) and DAPI for 30min. Stained samples were analyzed on an LSR II flow cytometer (BD) and cell numbers normalized using counting beads (Thermo).

### CodePlex secretome assay

Supernatants were harvested from overnight co-cultures and stored at −80°C until further use. On the day of the assay, samples were thawed and 11μl of supernatant per sample was added to CodePlex Human Adaptive Immune secretome chips (Isoplexis). Chips were loaded into the Isolight reader and cytokines measured using default settings. Automated analysis of raw data was performed using IsoSpeak software (Isoplexis).

### Statistical Analysis

Significance of differences in cell numbers, cytokine levels, and mean fluorescence intensity levels were calculated by Student’s t-test. All statistical tests were performed using Prism 9 (GraphPad Software). Results were considered significant when *p* <0.05.

### Study approval

The retrospective analysis of patient samples for the expression of CD229 was approved by the Institutional Review Board at the University of Utah (protocol 77285-19). Animal experiments were approved by the institutional animal care and use committee at University of Maryland Baltimore (protocol: 1021001).

## AUTHOR CONTRIBUTIONS

EVM, and TL conceived the project, planned and performed experiments, analyzed data, and wrote the manuscript. JMB performed variant scFv expression and CAR T cell production, and analyzed data. PD and DPN performed flow cytometry analysis of MM samples, and analyzed data. KD performed flow cytometry analyses and analyzed data. DA, SVR, DN, JM, MS, MLO, and CL analyzed data. All authors reviewed and approved the manuscript.

## ACKNOWLEDGEMENTS

This project was supported by a NCCN Young Investigator Award (T.L.), Huntsman Cancer Institute Experimental Therapeutics program supported by the National Cancer Institute of the National Institutes of Health under Award number P30CA042014. (D.A. and T.L.). We thank the Huntsman Cancer Institute in Salt Lake City, UT for the use of the Preclinical Research Resource (PRR), which performed the MM xenograft model, the ARUP Institute for Clinical and Experimental Pathology, and the University of Utah Flow Cytometry core facility, which assisted with flow cytometry analyses and cell sorting. pHIV-Luc-ZsGreen was a gift from Bryan Welm (Addgene #39196), SFG.CNb30_opt.IRES.eGFP was a gift from Martin Pule (Addgene # 22493), pSANG10-3F (Addgene plasmid # 39264) was a gift from John McCafferty.

