## Supplementary Material for "Combinatorial T cell engineering eliminates on-target off-tumor toxicity of CD229 CAR T cells while maintaining anti-tumor activity"

**SUPPLEMENTARY FIGURES**

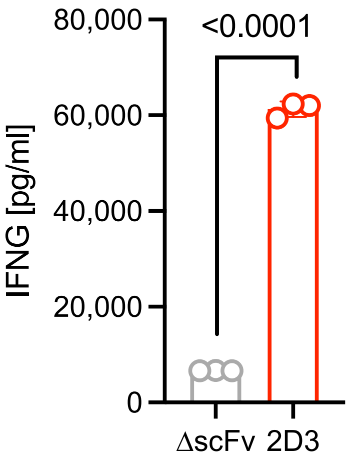


**Supplementary Figure 1: IFNγ production by CD229 CAR T cells co-cultured with U266B1 cells.** CAR T cells were co-cultured overnight with U266B1 cells at an effector-target ratio of 1:1. Supernatants were harvested and analyzed by IFNγ ELISA. Data indicate mean ± S.D. from technical replicates (*N*=3). Statistical significance was determined by two-tailed Student's *t* test.

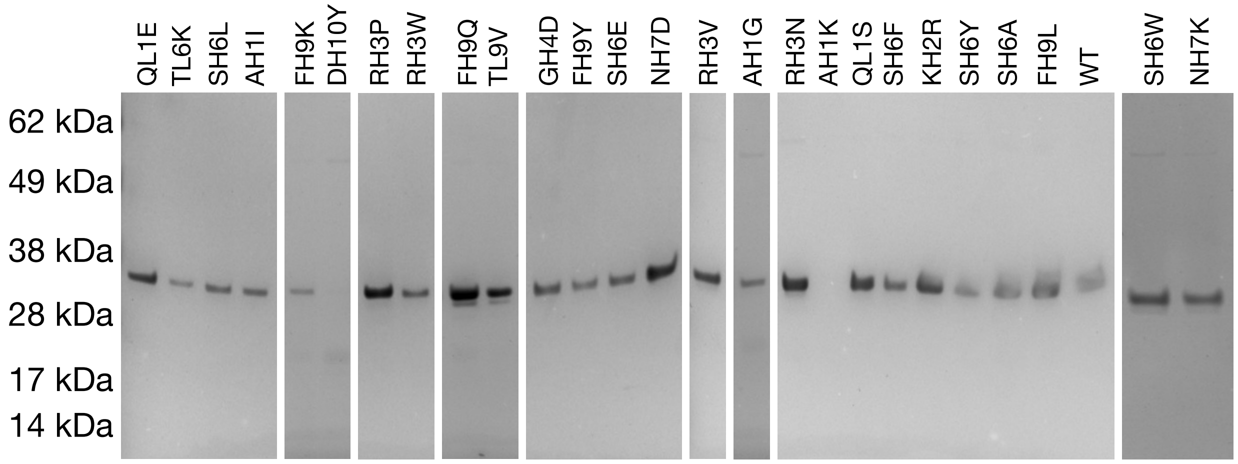


**Supplementary Figure 2: Purity of biotinylated 2D3 variants.** A fixed volume of each *in vivo* biotinylated antibodies was subjected to SDS-PAGE immediately after NiNTA purification and dialysis. Gels were stained with GelCode Blue (Thermo) and imaged using an iBright imaging system (Thermo).

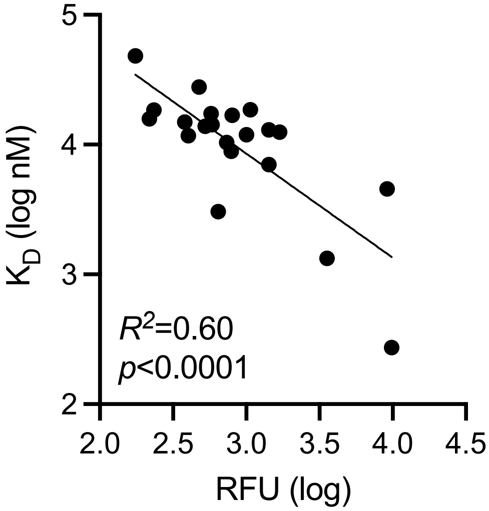


**Supplementary Figure 3: Correlation of variant binding by solid phase TRF assay and equilibrium constant.** For the solid phase assay 2ng/ul 2D3 variant scFvs were incubated with immobilized recombinant CD229 and binding was determined using anti-FLAG (clone: L5) and anti-mouse IgG-Eu (Perkin-Elmer). Equilibrium constants were determined by BLI. Significance of correlation was determined by Pearson *r* test and two-tailed *p* value.

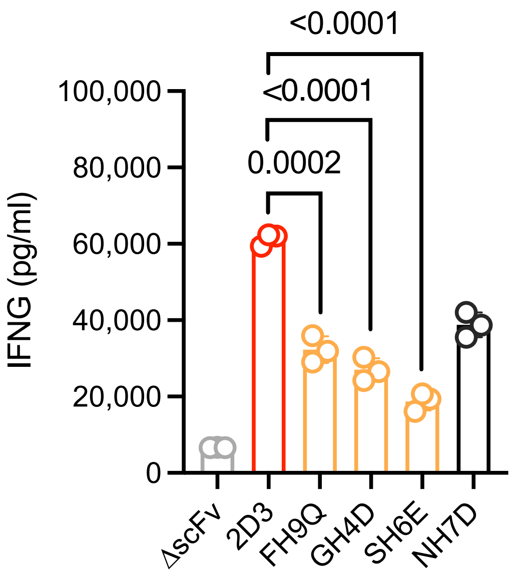


**Supplementary Figure 4: IFNγ production by CD229 variant CAR T cells.** CAR T cells were co-cultured overnight with U266B1 cells at an effector-target ratio of 1:1. Supernatants were harvested and analyzed by IFNγ ELISA**.** Data indicate mean ± S.D. from technical replicates (*N*=3). Statistical significance was determined by two-tailed Student's *t* test.

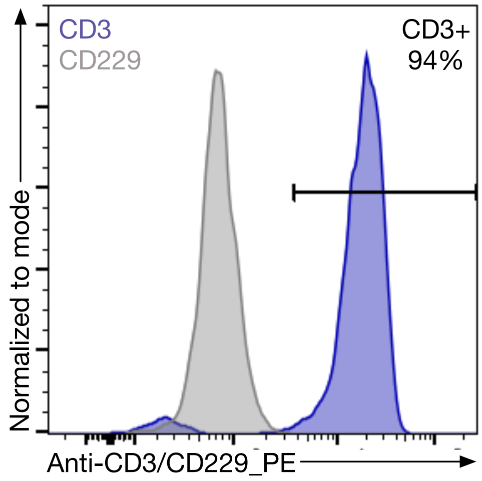


**Supplementary Figure 5: Purity of target T cells following negative selection.** Healthy T cells were purified by negative selection (Stem Cell Technologies), stained with anti-CD3/PE or anti-CD229/PE antibodies, and purity determined by flow cytometry.

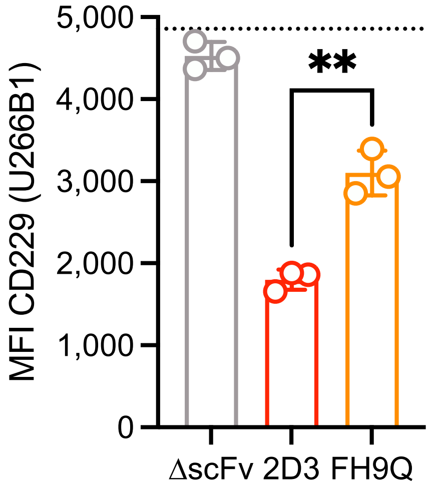


**Supplementary Figure 6: CD229 loss from U266B1 cells after CD229 variant CAR T cell co-culture.** U266B1 cells were labeled with CellTrace Far Red and incubated with CD229 variant CAR T cells for 4 hours. Co-cultures were stained with an anti-CD229/PE antibody and CD229 MFIs on CellTrace-positive U266B1 cells determined by flow cytometry. Data indicate mean ± S.D. from technical replicates (*N*=3). Statistical significance was determined by two-tailed Student's *t* test.

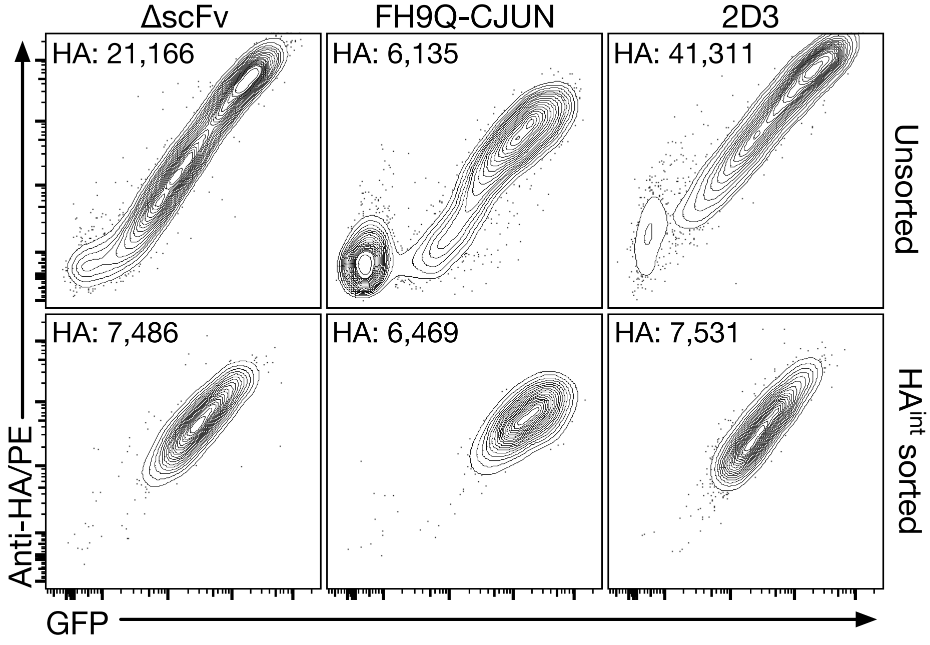


**Supplementary Figure 7: CAR surface expression levels before and after cell sorting.** Following CAR T cell production, cells were stained with an anti-HA antibody to determine CAR surface expression levels and subsequently sorted by flow cytometry. CAR and GFP expression were determined in sorted and unsorted cell products following anti-HA staining by flow cytometry. Mean fluorescence intensity for anti-HA-PE of the shown population is indicated in each panel.

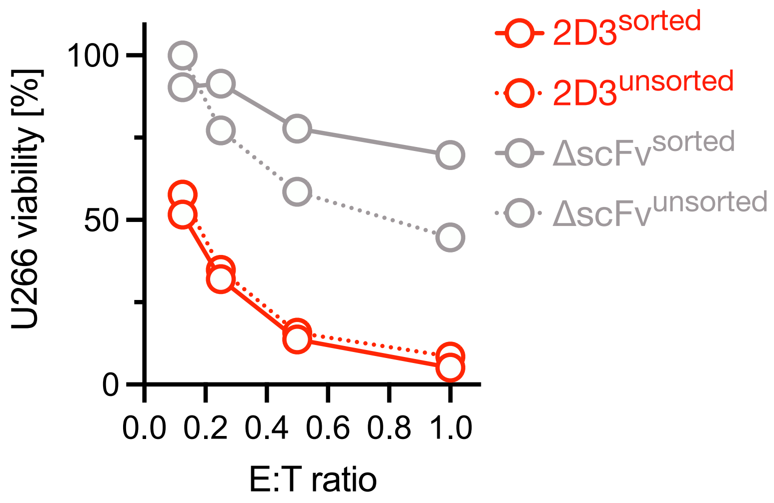


**Supplementary Figure 8: Killing of MM cells by CD229 CAR T cells following normalization of CAR surface expression levels.** Using a luminescence-based cytotoxicity assay, killing of luciferase-expressing U266 cells by CD229 CAR T cells before and after sorting for comparable HA/CAR expression was determined. Data indicate mean ± S.D. from technical replicates (*N*=3). Statistical significance was determined by two-tailed Student's *t* test.

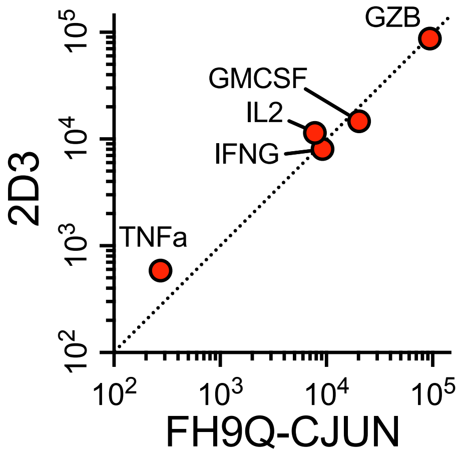


**Supplementary Figure 9: Cytokines secreted by CD229 CAR T cells during co-culture with MM cells.** FH9Q- and 2D3-based CAR T cells were incubated with U266 cell over night at an effector-target ratio of 0.5:1 and supernatants subjected to Isoplexis CodePlex analysis. Data indicate mean from 2 technical replicates.

**SUPPLEMENTARY TABLES**

| Target | Clone | Fluorophore | Supplier | Target cells |
|---|---|---|---|---|
| huCD229 | HLy9.1.25 | PE | Biolegend/R&D Systems | Multiple |
| huCD3 | UCHT1 | BUV496 | Biolegend | Pan-T |
| huCD138 | MI15 | BV510 | Biolegend | MM/Plasma |
| icVS38c | vs38c | FITC | Dako/Agilent | Primary MM |
| CD56 | N901 | ECD | Beckman | Primary MM |
| icKappa | TB28-2 | APC | Biolegend | Primary MM |
| icLambda | MHL-38 | APC-A750 | Biolegend | Primary MM |
| CD38 | HB7 | BUV395 | Fisher | Primary MM |
| CD19 | HIB19 | BUV737 | BD | Primary MM |
| CD45 | HI30 | AF790 | Thermo | Primary MM |
| Hemagglutinin | 6E2 | PE/APC | Cell Signaling Technology | CAR T |
| LIVE/DEAD Aqua | L34957 | N/A | Life Technologies | Live/dead |
| Propidium iodide | 421301 | N/A | Biolegend | Live/dead |
| DAPI | D1306 | N/A | Life Technologies | Live/dead |
| DYKDDDDK | M2 | PE | Sigma-Aldrich | Multiple |
| DYKDDDDK | L2 | PE/APC | Biolegend | Multiple |

**Supplementary Table 1: Table of monoclonal antibodies and viability dyes used for flow cytometry analyses.**
